# How the Tulip Breaking Virus Creates Striped Tulips

**DOI:** 10.1101/2024.06.05.597607

**Authors:** Aidan Wong, Gustavo Carrero, Thomas Hillen

## Abstract

The beauty of tulips has enchanted mankind for centuries. The striped variety has attracted particular attention for its intricate and unpredictable patterns. A good understanding of the mechanism that drives the striped pattern formation of the *broken tulips* has been missing since the 17th century. It is known since 1928 that these patterned tulips suffer from a viral infection by the *tulip breaking virus*. Here, we present a mathematical model to understand how a virus infection of the petals can lead to stripes, thereby solving a 350 year old mystery. The model, which describes the viral inhibition of pigment expression (anthocyanins) and their interaction with viral reproduction, incorporates a pattern formation mechanism identified as an *activator-substrate* mechanism, similar to the well-known Turing instability, working together with a Wolpert’s positional information mechanism. The model is solved on a growing tulip petal shaped domain, whereby we introduce a new method to describe the tulip petal growth explicitly. This work contributes to the theory of pattern formation of reaction-diffusion systems on growing domains applied to the fields of virology and botany.

---

Leaves, petals, and plants are full of vibrant colours and beautiful patterns. Gardeners try their best to defend their plants from botanical diseases. Few would think that viruses can create beauty in our world. However, tulips infected with the tulip breaking virus (TBV) can generate petals with streaks, stripes, or flames (see Figures 1a-b).

**Figure 1:**
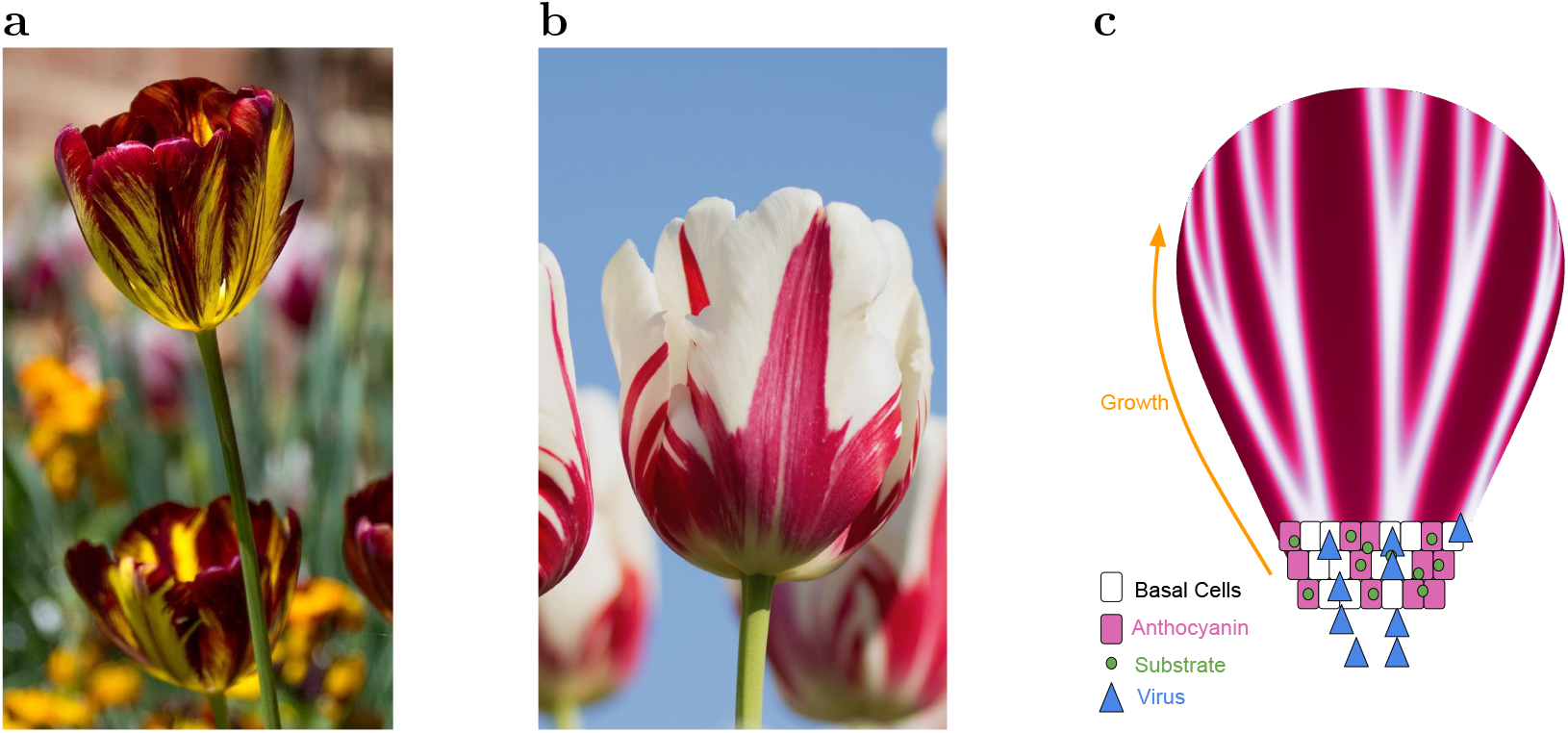
**a**, *Tulipa Absalon* variety, one of the few maintained “truly” broken bulbs that is striped from the tulip breaking virus rather than genetics. Yellow denotes a loss of pigments, and suggests those regions are infected. Reproduced with permission from The Mount Vernon Ladies’ Association ^32^. **b**, Tulip of the variety *Carnaval de Rio*. Image taken by Kwang Mathurosemontri, available on `unsplash`. **c**, A schematic of the components of the TBV model (1-3). Along the base of the petal are layers of cells, each with varying concentrations of anthocyanin pigment due to the virus consuming the substrate and silencing pigment expression.

Tulips were brought from Turkey to the Netherlands in the 1590’s by the ambassador Ogier Ghislain de Busbecq^17^. The Dutch botanist Carolus Clusius is credited with having planted the first tulips in Holland. They were so popular that his garden was frequently robbed until he eventually gave away his collection, spreading tulips throughout the Netherlands ^17^. During his studies, Clusius noticed that some tulips had striped patterns, but these were also smaller, weaker, and less likely to reproduce. It seemed that their beauty had come with frailty, and he suggested they were diseased ^17^.

The beauty of these broken bulbs enchanted the Dutch population during the 17th century, causing a soar in tulip prices and a subsequent plummet. This phenomenon is referred to as Tulipomania, and some economists allude to it as the first recorded financial bubble^23;33^. During the Tulipomania, it was recognized that colour breaking was transmissible from broken tulips to healthy, solid tulips by aphids. The first identified virus, the Tobacco Mosaic Virus (TMV), was discovered in 1892^38^. Only later, in 1928, could Cayley and McKay^8^ identify the virus that is responsible for the broken tulip patterns, the *tulip breaking virus* (TBV). This fascinating history of broken tulips is gracefully portrayed by Deborah Leipziger in her poem *The Broken Tulip*^14^:

*For decades, no one knew*

*what caused the flaring*

*the feathering of tulips*,

*Parrot like*,

*Red on orange*

*Peppermint red on white*

*Black on tangerine —*

*The eruption into flame*

*for broken tulips like*

*Absalom and Mabel*

*What causes tulips to “break”*

*The mosaic virus*

*carried by aphids*

*infects bulbs*

*and the flower breaks*

*its hold on one color*,

*the primary color suppressed*

*and lighter colors bleeding through*

*the beauty of a curse*

Despite the knowledge of the TBV, the process of how a virus infection of the petal can lead to spatial pattern formation remains to be understood. Karin Moelling ^24^ formulated this as an open question in her book from 2017. She writes on pages 223-224: ”*Alan Turing* … *described the mathematics leading to stripes*. … *an activator and a long-range inhibitor are interacting - but I do not know which of the two is the virus*.”

The answer to Dr. Moelling’s question is a bit more complicated than expected. We shall see that the pattern forming mechanism in broken tulips is that of an *activator-substrate model* (also called *activator-depletion model*)^10^. Unlike an activator-inhibitor system, we have a long-range substrate instead of having a long range inhibitor. In that context, *the virus is the activator*, which inhibits anthocyanin expression.

Although patterns are prevalent everywhere in biology, from the stripes on a zebra to the structure of fingers, their root factors are only understood in very limited cases. In 1952, the famous computer scientist and mathematician Alan Turing asserted that some patterns could be caused by chemicals that react and diffuse across a spatial domain^20;35^. Proposed reaction-diffusion equations have been successful in mimicking seashells^20^, fish stripes^26^, and mammalian skin patterns^25^. In 1977, Wolpert et al. proposed a simple gene activation model that could produce distinct thresholds for patterns in a diffusive environment^18^. In their model, cells retain positional information by sensing the concentration level along a gradient. From then on, these two theories have been competing with each other. However, given the great variety of biological functions, Turing and Wolpert’s theories have been shown to act in concert in many cases, to produce biological patterns^11;21^.

Considering animal skin patterns, an additional challenge arose. While an animal grows, its skin surface also grows and changes, and this often occurs on the same time scale as the pattern forming process. Edmund Crampin and collaborators developed methods to apply pattern forming models on domains that change over time^6^. Their method transforms a system of reaction-diffusion equations on a one-dimensional growing domain to a modified system on a fixed domain. Our model for the TBV will be based on both the Wolpert-type gradient sensing and the Turing instability mechanisms on a growing domain.

Turing instability has already been applied to model the formation of patterns in plants; in particular, to describe the spot formation in the flowers of Monkeyflowers (*Mimulus*)^7;37^. Monkeyflowers contain spots on their nectar guides, which consist of high levels of anthocyanin. This example gives great credence to using reaction-diffusion equations to model patterns in plants.

Here, we present a mathematical model that provides a non-linear dynamics explanation for the formation of petal patterns in broken tulips. The model is based on the interplay of virus infection with cell pigmentation gene expressions for *anthocyanin* and comprises a system of partial differential equations (PDEs), balancing the effects of viral infection, viral use of cell resources, and inhibition of gene expressions for cell pigments. The PDE system includes two famous pattern forming mechanisms, a Wolpert mechanism of pattern formation in spatial gradients ^18^, combined with a Turing instability mechanism^35^. These dynamics are embedded in a growing domain of a tulip petal. We use the methods developed by Crampin et al. ^6^ to introduce a growing domain into the PDE formulation, and develop a parametrization of a growing tulip petal to predict the formation of stripe patterns in broken tulips.

## Building the TBV model

To develop the model, we describe the spatio-temporal dynamics and biological reactions between three components of an infected tulip petal, namely the pigment, the virus, and the viral resources. The natural pigments in the tulip are represented with the concentration of anthocyanin, *T* (*x, t*), the primary pigment in red, black, and purple tulips^36^. The infection is induced by TBV viral components, measured in viral load *V* (*x, t*), that diffuse across plant cells filled with substrate, whose concentration is described by our last variable, *B*(*x, t*). As ”building blocks”, it represents resources inside cells such as proteins, amino acids, hormones, and nutrients that are used to build new virions. A similar substrate approach is found in Moreno’s work ^2^.

Assuming that these components diffuse across a spatial domain [0, *L*(*t*)], which grows in time as the plant grows, and that reactions are not dependent on location, temperature, or soil acidity, the model describing the dynamics of anthocyanin *T* (*x, t*), the virus load *V* (*x, t*), and the substrate *B*(*x, t*), is given as

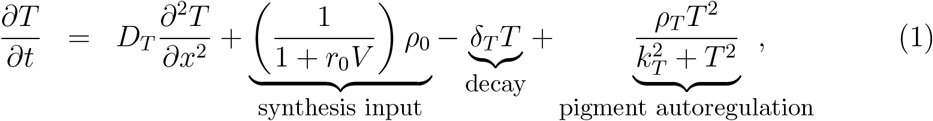

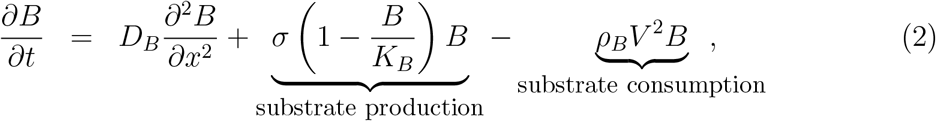

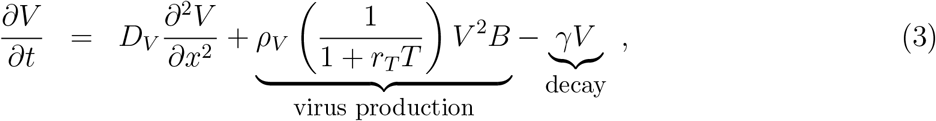

on a domain [0, *L*(*t*)] with homogeneous Neumann boundary conditions (see Supplement 2). The parameters of the model are defined in Table 1 within Supplement 1.

**Table 1:**
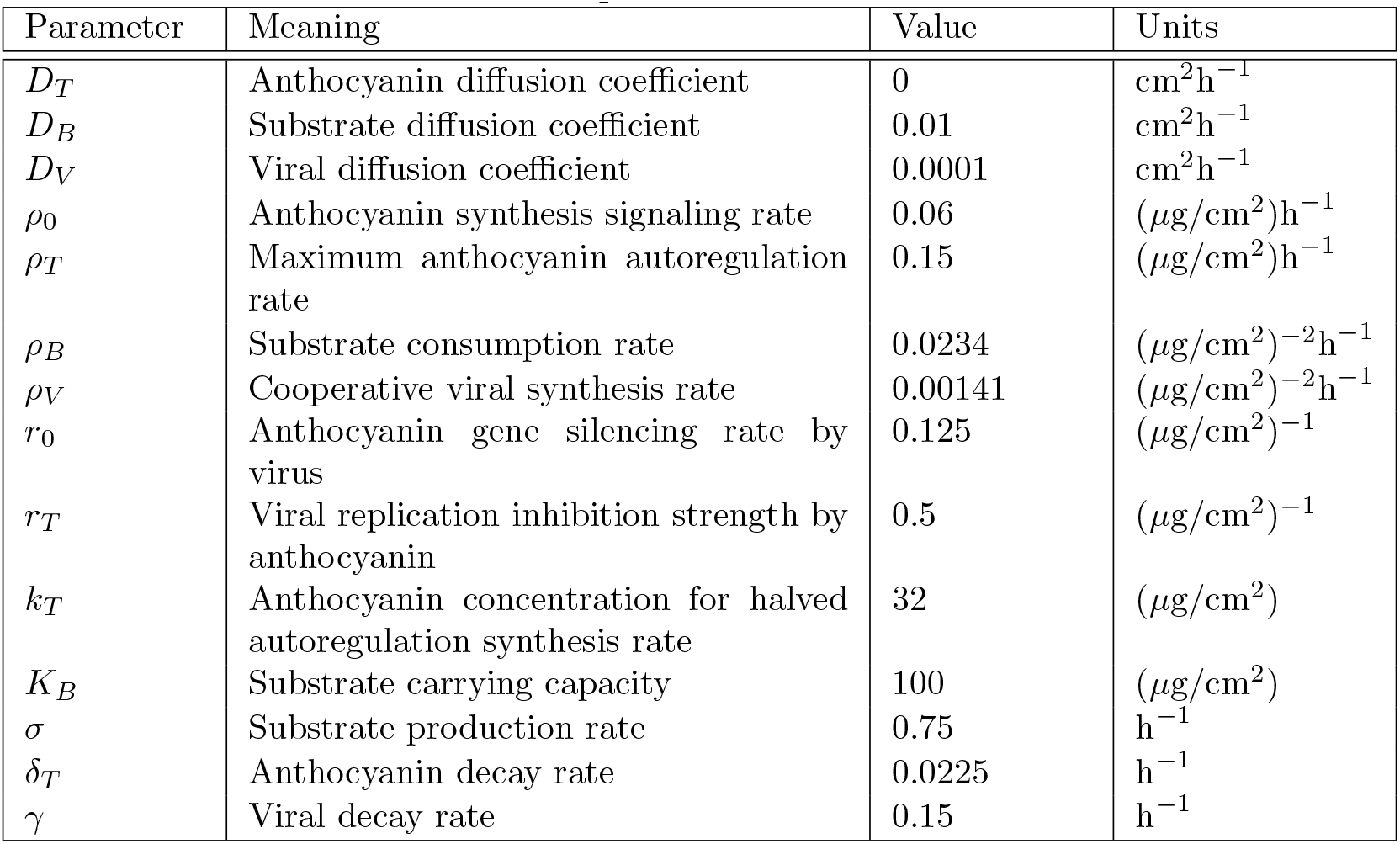
Dimensionalized parameter values for the TBV model.

| Parameter | Meaning | Value | Units |
| --- | --- | --- | --- |
| $D_T$ | Anthocyanin diffusion coefficient | 0 | $\text{cm}^2\text{h}^{-1}$ |
| $D_B$ | Substrate diffusion coefficient | 0.01 | $\text{cm}^2\text{h}^{-1}$ |
| $D_V$ | Viral diffusion coefficient | 0.0001 | $\text{cm}^2\text{h}^{-1}$ |
| $\rho_0$ | Anthocyanin synthesis signaling rate | 0.06 | $(\mu\text{g}/\text{cm}^2)\text{h}^{-1}$ |
| $\rho_T$ | Maximum anthocyanin autoregulation rate | 0.15 | $(\mu\text{g}/\text{cm}^2)\text{h}^{-1}$ |
| $\rho_B$ | Substrate consumption rate | 0.0234 | $(\mu\text{g}/\text{cm}^2)^{-2}\text{h}^{-1}$ |
| $\rho_V$ | Cooperative viral synthesis rate | 0.00141 | $(\mu\text{g}/\text{cm}^2)^{-2}\text{h}^{-1}$ |
| $r_0$ | Anthocyanin gene silencing rate by virus | 0.125 | $(\mu\text{g}/\text{cm}^2)^{-1}$ |
| $r_T$ | Viral replication inhibition strength by anthocyanin | 0.5 | $(\mu\text{g}/\text{cm}^2)^{-1}$ |
| $k_T$ | Anthocyanin concentration for halved autoregulation synthesis rate | 32 | $(\mu\text{g}/\text{cm}^2)$ |
| $K_B$ | Substrate carrying capacity | 100 | $(\mu\text{g}/\text{cm}^2)$ |
| $\sigma$ | Substrate production rate | 0.75 | $\text{h}^{-1}$ |
| $\delta_T$ | Anthocyanin decay rate | 0.0225 | $\text{h}^{-1}$ |
| $\gamma$ | Viral decay rate | 0.15 | $\text{h}^{-1}$ |

Let us briefly describe each of the terms in model (1-3). The production and decay of the anthocyanin *T* in equation (1) is modelled using the well-known Wolpert model^18^. It is a simple yet accurate model describing the production of a chemical via DNA synthesis in cells. The parameter *ρ*_0_ denotes a constant signal strength that activates the genes for *T*. Since strong evidence suggests that Potyvirus infection can lead to gene silencing in plants^5;29;1;12^, the activation *ρ*_0_ is reduced by the factor (1 + *r*_0_*V*)^−1^. The parameter *δ*_*T*_ denotes the natural degradation of anthocyanin. As there is evidence of an autoregulatory feedback loop of transcription factors responsible for the regulation of anthocyanin biosynthesis^9;27;19;34^, we describe a simple autoregulation mechanism for anthocyanin in the last term of equation (1), where *ρ*_*T*_ denotes the rate of growth that is halved when *T* = *k*_*T*_. In the absence of the virus (*V* = 0), the tulip naturally regulates the anthocyanin concentration onto a homogeneous and constant homeostatic value.

For the substrate equation (2), we use a simple logistic growth model with growth rate *σ* and carrying capacity *K*_*B*_. Since viral replication is proportional to the substrate consumption and cooperative in nature ^2;16;30^, the viral cooperative interaction appears in both the substrate consumption term of equation (2) and the virus production term of equation equation (3). This latter term is reduced by a factor (1 + *r*_*T*_ *T*)^−1^ to reflect the anti-viral properties of anthocyanin^17;12;1;31^. The virus is naturally degraded at a rate *γ*. Finally, all equations are equipped with diffusion terms. A schematic of the components of the TBV model is depicted in Figure 1c.

A comprenhensive choice of parameter values for the TBV model is described in the Supplement 1. Also, we nondimensionalize the model in Supplement 2; this allows us to reduce the number of parameters and use the nondimensional model to carry out a multiscale analysis and the numerical simulations.

### Incorporating the tulip petal growth into the model

The base of the flower petal can be identified as a one-dimensional interval, [0, *L*(*t*)], that lengthens over time and creates layers of a petal that continues to grow during a TBV infection. In order to examine how the growth and curvature of tulip petals evolve, we take a closer look of a tulip petal in Figure 2a, which shows its fan-like veinal structure. This structure leads us to assume that cells aligned with the same perpendicular arc to the veins have the same age.

**Figure 2:**
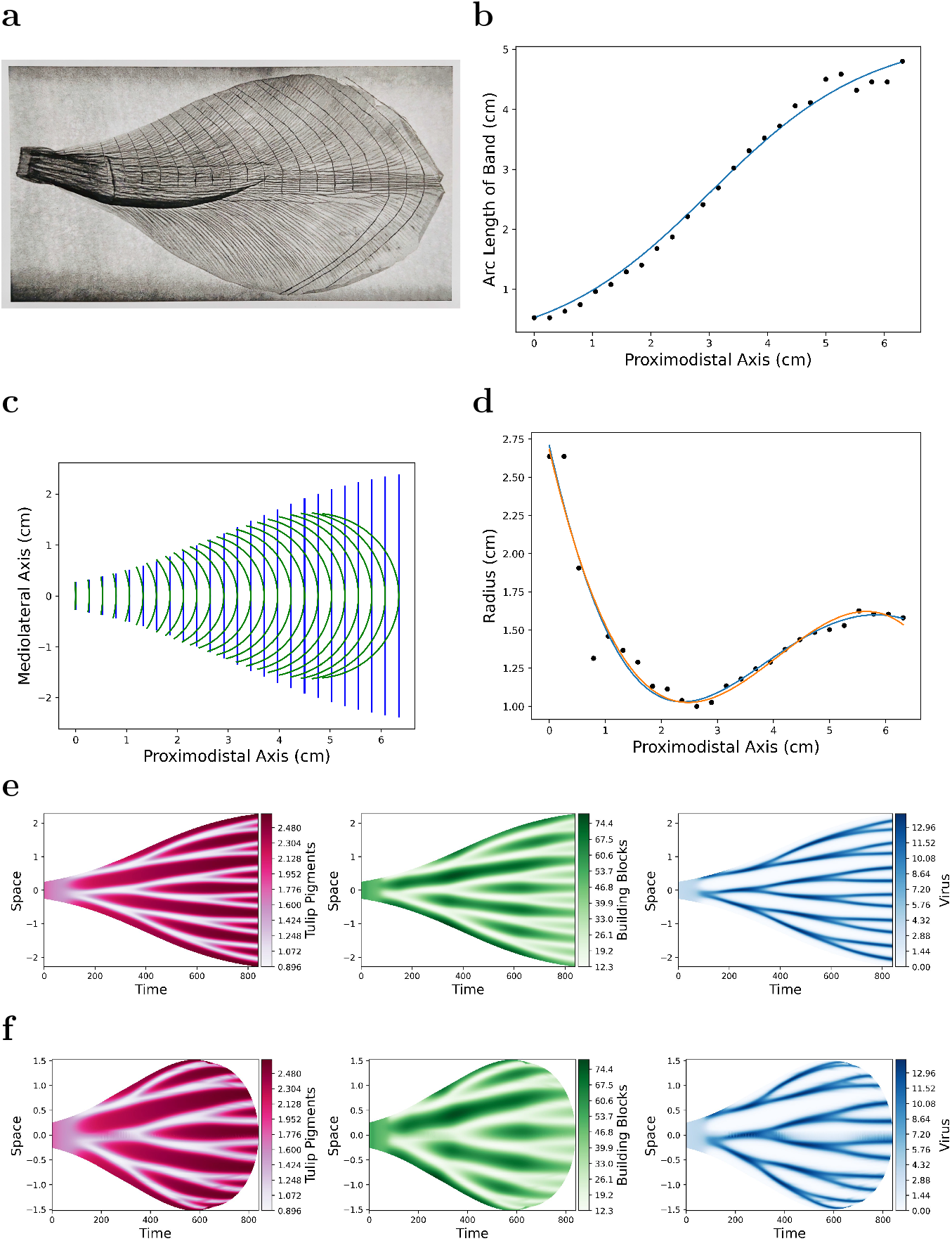
The growth and curvature of tulip petals. **a**, A visible light image of a petal from *“Tulipa Seadov”* printed on paper. Lines were drawn from evenly spaced points on the 6.35 cm proximodistal (base to tip) axis, and then perpendicular to each vein. **b**, Estimated circular arc lengths of the perpendicular arcs to the petal veins with respect to their position on the proximodistal axis. The fitted logistic curve *L*(*t*), given by Equation 4. **c**, The transformation of the intervals [0, *L*(*t*)] from straight blue lines into green arcs according to the circular parametrization. **d**, Radii of the circular arcs in figure c. A Wolpert function (blue) and a cubic polynomial (orange) were fitted. **e**, Dynamics of the TBV model on the domain [0, *L*(*t*)] with logistic growth given by Equation (4) over 35 days, with parameters *L*_0_ = 0.5 *cm, κ* = 0.727, *ξ* = 9.936 and *τ* = (35 · 24 h)/6.35 = 132.28 h. Time is given in hours and space is in cm. **f**, Dynamics of the TBV model on the corresponding parametrized arcs (curved green circular arcs in figure e with radii *R*(*t*) = −0.037205(*t*/*τ*)^3^ + 0.454(*t*/*τ*)^2^ −1.565(*t*/*τ*) + 2.682, with the same value for *τ*). Time applies to the intersection of each circular arc with the center line.

Even though tulip petals come in a wide array of shapes and sizes, we choose to illustrate, in Figure 2a, a tulip petal of the variety *“Tulipa Seadov”*, and use this triumph tulip as a representative sample. Perpendicular arcs to the petal veins are drawn equidistant along the proximodistal (base to tip) axis of the petal. These arcs are parametrized, for the purpose of generating the simplest growing tulip petal shaped domain, as circular arcs (see Supplement 5 for details). This parametrization allows us to estimate the circular arc lengths, illustrated in Figure 2b, where the *x*-axis, denoting the distance along the proximodistal axis, is associated with the growth time.

The estimated arc length grows logistically and thus fitted with the function

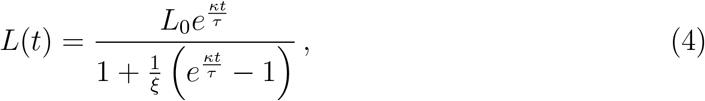

with parameters *κ* and *ξ*, and timescale *τ*, determined by the time it takes for the tulip flower to start growing and mature. The typical time for this is 30 to 40 days^22 15^. Assuming that the growth is homogeneous and knowing the length of the proximodistal axis of a sample petal is 6.35 cm, we can create functions of petal growth with respect to time by relating the proximodistal length to the flowering time. These functions return arc lengths that correspond to the lengths of the growing domain [0, *L*(*t*)] for our TBV model. This correspondence, shown in Figure 2c, is obtained by estimating the arc radii. Figure 2d displays these radii values and their non-monotonic behaviour fitted with a Wolpert and a cubic polynomial function (see Supplement 5 for more details).

### Emergent stripe patterns on a growing tulip petal

Armed with the analytic function (4) describing the domain growth, we can now solve the TBV model (1-3) on a growing tulip shaped domain. For this purpose, we implement in the TBV model (1-3), the procedure developed by Crampin et al. ^6^ to transform a reaction-diffusion model on a growing domain [0, *L*(*t*)] into a modified model on a fixed domain (see Supplement 4 for details).

To illustrate the results, we start with an initial state where concentration of viral components is randomly distributed with a standard deviation of 0.1 around its kinetic steady state. We let the dynamics of the TBV model evolve first on the growing domain of length *L*(*t*), indicated as straight blue lines in Figure 2c, which results in the emergence of spatial patterns of its components shown in Figure 2e. Thereupon, we identify these spatially patterned dynamics onto the corresponding parametrized arcs (curved green lines in Figure 2e), and obtain the stripe patterns on a tulip shaped petal domain, shown in Figure 2f.

#### The Forward and Backward Models

The simulations carried out so far, which start with an initial condition at the petal base, presuppose that all the new layers of the tulip petal emerging from its base share the same initial state. We call this case, the *Forward Model*. We can also carry out simulations starting with an initial condition at the outer rim, and consider the case in which a new layer of the TBV model components emerging from the petal base is assumed to be solely influenced by the previous layer. This case is named the *Backward Model*. Note that even though the simulations for the two models have opposite initial domains (one at the base and the other at the tip of the petal), the oldest part of the tulip petal is the outer rim in both cases.

In Figure 3, we arranged six simulated tulip petals infected by TBV on different domain length sizes for the two above mentioned models, and notice fewer stripes in smaller domains and more stripes in larger domains. This is expected for an activator-substrate mechanism. Another striking result comes from the pattern formation differences between the two models. Observing the region close to the outer rim of the petals, we notice that for the *forward* case the effect of the viral pattern formation dynamics kicks in as soon as the petal starts growing (petals in Figure 3a), whereas the effect of the virus on the formation of patterns is felt by the petal some time after for the *backward* case, resulting in a transient period without any patterns (petals in Figure 3c). Interestingly, this difference in tulip petal patterns was captured during the Dutch Golden Age by artists who wished to preserve the beauty of broken tulips in their paintings (Figures 3b and 3d).

**Figure 3:**
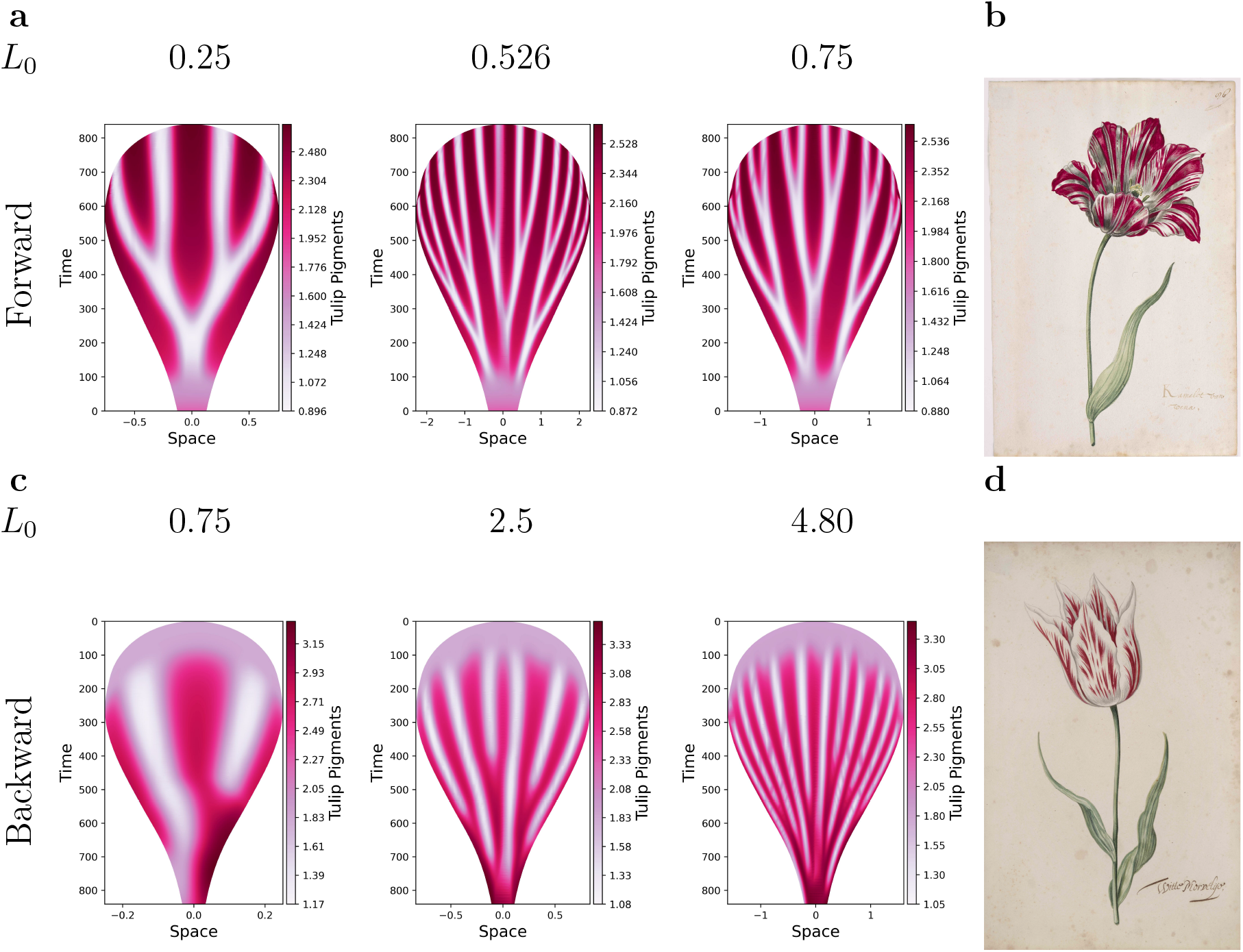
Simulations of tulip petals infected by TBV. Initial conditions are the model’s kinetic steady state concentrations except for the virus concentration, which is randomly distributed with a standard deviation of 0.1 around its kinetic steady state. **a**, Dynamics of the TBV model on tulip petals with varying initial base lengths that grow forwards and towards the broad rim. This *forward* dynamic assumes that all the new layers of the tulip petal emerging from its base share the same initial state. The initial length *L*_0_ is listed above each petal in centimetres. *L*_0_ = 0.526 is realistic for forwards growth, while *L*_0_ = 4.80 is realistic for backwards growth. **b** The tulip *Kamelot van Wena*, painted by an unknown Dutch Artist in the 17th century. Reproduced with permission from Norton Simon Art Foundation ^3^. **c**, Dynamics of the TBV model on tulip petals with varying initial outer rim lengths that grow backwards and towards the petal base. This *backward* dynamics assumes that new layers of the TBV model components emerging from the petal base is solely influenced by the previous layer. **d**, The tulip *Witte Merveljie*, painted by an unknown Dutch Artist in the 17th century. Reproduced with permission from Norton Simon Art Foundation ^4^. There are intriguing similarities between the petals of the paintings in figures b and d and the simulated petals of figures a and c, respectively.

### Turing and Wolpert mechanisms working in unison explain stripe patterns

A thorough multiscale and Turing stability analysis of the TBV model (1-3) is carried out in Supplement 3. Based on our comprehensive choice of parameter values, it comes to light that the substrate and the virus act on a faster time scale than the pigmentation. The substrate-virus sub-system is responsible for the formation of patterns, and we can say they form a pre-pattern. The slower *T* equation then amplifies this pre-pattern by expressing high pigment levels for low virus concentrations and low pigment levels for locations of high virus load. In other words, the Turing model reactions are created upstream of the positional information^11^.

## Discussion

We set out to solve a flower mystery that has served as a source of inspiration for artists and writers since the 17th century, the delightful stripe patterns of *broken tulips*. This mystery started to reveal itself in 1926, when it was discovered that the stripes are caused by the *tulip breaking virus* infection of the tulip petals. However, how this infection leads to stripe patterns has been an open question ever since.

In this article, we develop a mathematical model to explain the underlying instability that leads to stripe patterns, thereby answering the above open question and contributing to the full unfolding of this 350 year old mystery.

The TBV model (1-3) identifies the dynamic interaction between the tulip breaking virus and its resources as a substrate-virus system responsible for the formation of Turing-like patterns and taking place at a faster time scale than the pigment dynamics, which is described by a Wolpert positional information process. This interplay between these two pattern formation mechanisms (Turing’s and Wolpert’s) acting together on different time scales, and embedded on a growing petal shape domain parametrized with respect to the veinal structure growth of tulip petals, results in the emergence of stunning stripe patterns.

In addition to portraying the beauty of broken tulips within a pattern formation frame built with the robust theories of Turing and Wolpert working in unison, we have expanded upon and applied Crampin’s method of modelling growing domains to a real, biological model, namely the TBV model. The model assumes that a petal grows sequentially, layer on layer of cells, and therefore taken as a one dimensional domain that grows uniformly along its axis. More applications of growing domains have been long overdue because many biological surfaces are not static. Hopefully this paper can reveal the potential of this powerful method, that may extend to more biological systems, such as the stripes generated by genetically bred tulips.

We assume that TBV silences the gene that expresses anthocyanin rather than interfering with its synthesis or regulation. While viral gene silencing has been observed^5;29;1;12^, it is not the definite, or the only possible process^17^.

Nowadays, ”true” broken bulbs infected with viruses are not available for popular distribution. Fortunately, for fans of their beauty, natural striped tulip varieties exist. It is uncertain if the TBV model can be applied to engineered tulip varieties. One can make a case that transgenes^13^, MYB proteins^36^, other proteins, or a combination of them can serve as a surrogate breaking virus within natural varieties since they can suppress genes and interfere with pigment regulation. However, natural regulators should not consume the substrate at a rate we expect from viruses.

Finally, the beauty of the patterns generated by the TBV model is astonishing. We could generate a wide variety of artworks by simply changing few parameters and colours. Most Turing patterns are spots or hatches, but branching stripes can be another addition to the repertoire of mathematical art.

## Acknowledgements

TH is grateful to C. Friedhoff for exploration of her Tulip garden. TH research is supported by an NSERC Discovery grant RGPIN-2023-04269. AW and GC acknowledge the support received from the Athabasca University ARF (Academic Research Fund) # 25236.

## Author Contributions

TH conceptualized the study. GC designed the model. AW contributed to the programming. All authors contributed to the final version of the manuscript.

## Competing Interests

The authors declare no competing interests.

## Additional Information

**Supplementary Information** is available for this paper.

**Peer review information** *Nature* thanks the anonymous reviewers for their contribution to the peer review of this work. Peer reviewer reports are available.

## Supplementary Information

### 1. Supplement 1: Choice of parameter values

In Table 1 we summarize the parameter values that are used in our model. Since there are few specific studies on TBV, but more on the similar rod-like virus TMV, sources on TMV are often consulted.

First, the average area of a petal ranges from 13 to 25 cm^2^, and the fresh weight of a petal ranges from 0.5 to 0.9 g ^2^. Given this range, the density of petal mass per area is approximately 0.04 g cm^−2^. Most values in the literature measure the concentration of substances with (*µ*g/g), so we must multiply this density factor with values dependent on the weight *g* to retrieve units of concentration with respect to space rather than mass.

(*D*_*T*_) Anthocyanin does not diffuse freely within nor between petal cells. Rather, they are transported upon their synthesis into vacuoles for storage, from which we can see a flower’s bright colours. In other words, anthocyanin tends to stay where it is produced, and thus its diffusion coefficient is *D*_*T*_ = 0 ^24^.

(*D*_*B*_) The diffusion coefficient of auxin, (a vital plant hormone) within root cell cytoplasm is 220 *µ*m^2^/s, a third of its aqueous value^18;19^. The diffusion coefficient of glucose within immobile clusters of plant cells is 0.028 to 0.28 × 10^−5^ cm^2^/s ^15^. Additionally, the diffusion coefficient of sucrose in an aqueous solution is 0.521 × 10^−5^ cm^2^/s, with the diffusion coefficients of other sugars (lactose, fructose, glucose) similarily between 0.5−0.7 × 10^−5^ cm^2^/s ^22^. Finally, standard diffusion coefficients of water isotopologues within leaves range from 2.34 to 2.66 ×10^−5^ cm^2^/s at 25 ^°^C ^5^. All in all, the diffusion coefficient most likely ranges on an order of magnitude from 10^−6^ to 10^−5^ cm^2^/s. Hence, for the diffusion of the substrate *B* we use a value of *D*_*B*_ = 2.7×10^−6^ cm^2^/s, which is equivalent to *D*_*B*_ = 0.01 cm^2^/h.

(*D*_*V*_) While there seem to be no publications on the diffusion coefficient of TBV, there are some sources for the Tobacco Mosaic Virus (TMV), which, like TBV, is also a rod-shaped positive strand type RNA virus. Light scattering methods return values ranging from 3.75 to 4.50 × 10^−8^ cm^2^/s in buffer solutions^20;21^. Within a colloidal medium the value was 3.90 *µ*m^2^/s, and within a light suspension it was around 4.19 *µm*^2^*/s* ^12^. Also, the diffusion coefficient of viruses are typically around 1 % of the diffusion coefficient of the substrate it is in. Hence, we choose *D*_*V*_ = 2.7 × 10^−12^ m^2^/s, which is equivalent to *D*_*V*_ = 0.0001 cm^2^/h.

(*ρ*_0_, *ρ*_*T*_, *k*_*T*_, *δ*_*T*_) The coefficients for the Wolpert gene expression dynamics of anthocyanin were obtained by fitting the Wolpert model to data from various studies that measured the concentration of anthocyanin at several developmental stages of flowers^23;9^. In a study on transcription factors that synthesize and regulate anthocyanins, Tian et al. recorded the concentration of anthocyanin within crabapple flowers for five stages across 150 days^23^. Likewise, Guo et al. studied the concentration of anthocyanin with the Queen of Night tulips as the bud developed ^9^.

The Wolpert model returns 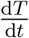(the change of the concentration of anthocyanin with respect to time) as a function of *T* (concentration of anthocyanin). See Figure 1 for a graph of the Wolpert model. Taking the data of anthocyanin concentration vs. time and calculating 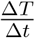, and then plotting it against the anthocyanin concentration gives us several points to fit with the Wolpert model with. The resulting parameters are *ρ*_0_ = 1.5 (*µ*g/g)h^−1^, *ρ*_*T*_ = 3.75 (*µ*g/g)h^−1^, *k*_*T*_ = 800 (*µ*g/g), and *δ*_*T*_ = 0.0225 h^−1^. After multiplying by our factor of petal mass density, we have *ρ*_0_ = 0.06 (*µ*g/cm^2^)h^−1^*ρ*_*T*_ = 0.15 (*µ*g/cm^2^)h^−1^, *k*_*T*_ = 32 (*µ*g/cm^2^), and *δ*_*T*_ = 0.0225 h^−1^.

(*ρ*_*B*_) This parameter denotes the cooperative consumption of the tulip’s proteins and nutrients by the virus. Its rate is derived from the work of Hagiladi et al.’s research into the Tobacco Mosaic Virus, which is discussed in more detail in the description of *ρ*_*V*_ below^10^. We assume that the substrate consumption rate *ρ*_*B*_ will be similar in magnitude to the viral production constant *ρ*_*V*_, since the substrate is subsumed into viral components. We have a range of values for *ρ*_*V*_ between 10^−7^ and 10^−5^ (*µ*g/g)^−2^h^−1^. We choose *ρ*_*B*_ = 3.75 × 10^−5^ (*µ*g/g)^−2^h^−1^. After multiplying our factor for petal mass density, we have *ρ*_*B*_ = 0.0234 (*µ*g/cm^2^)^−2^h^−1^.

(*ρ*_*V*_) The viral production constant was determined from the work of Hagiladi et al.’s research into the Tobacco Mosaic Virus. They studied the linear rate of change in concentration of TMV within leaves, and emerged with a value of 64.7 (*µ*g/g)h^−1^. The order of magnitude of TMV concentration within leaves is about 10^2^ (*µ*g/g) ^10^ and the order of magnitude of the substrate concentration is between 10^2^ to 10^3^ (*µ*g/g) ^6^. Reflecting on our model, we can approximate the linear rate as our viral production term in the absence of anthocyanin, as leaves have little or no concentration of it. Mathematically, this means 64.7 (*µ*g/g)h^−1^ *ρ*_*V*_ ≈ *V* ^2^*B*. Solving for *ρ*_*V*_ with our range of *V* and *B* gives us a range of values for *ρ*_*V*_ between 10^−7^ and 10^−5^ (*µ*g/g^2^)^−2^h^−1^. We chose *ρ*_*V*_ = 2.25×10^−6^ (*µ*g/g^2^)^−2^h^−1^. After applying our conversion factor for petal mass density, we have *ρ*_*V*_ = 0.00141 (*µ*g/cm^2^)^−2^h^−1^.

(*r*_0_) The gene signal inhibition strength was estimated in order to follow mathematical conditions for diffusion driven instability. After estimating a value for *r*_0_, we can solve for the nondimensional parameters *η* and *β*, and see if they follow the necessary conditions for diffusion driven instability. We can confirm that they produce patterns by running the simulation. Also, after integrating the concentration of anthocyanin and viral components over the 2D space of the petal, we expected realistic values within the same order of magnitude of the concentration of anthocyanin and viruses, which was achieved. A value of *r*_0_ = 0.125 (*µ*g/cm^2^)^−1^ was chosen.

(*r*_*T*_) Hayashi et al. ^11^ analyzed the ability for anthocyanin to inhibit human influenza viruses within aqueous solutions. They determined IC_50_ (the concentration of anthocyanin required to half the activity of this virus) to be 48 *µ*g/mL for influenza virus A and 54 *µ*g/mL for influenza virus B. Since the substrate is mostly an aqueous solution, and 1 mL of water weighs 1 g, we can do a rough unit change from (*µ*g/mL) to (*µ*g/g). Since the cooperative rate of viral growth is proportional (1 + *r*_*T*_ *T*)^−1^, it is halved when 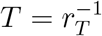. Since the IC_50_ is the concentration of anthocyanin at which the viral activity is halved, then *r*_*T*_ = (IC_50_)^−1^ which gives us an estimate around 0.02 (*µ*g/g)^−1^. After applying our conversion factor from concentration in petal mass to concentration in petal space we have *r*_*T*_ = 0.5 (*µ*g/cm^2^)^−1^.

(*K*_*B*_) The carrying capacity in our model is the maximum concentration of the substrate. The total concentration of proteins in the perianth (the flower organs of the plant) of a tulip at full bloom was 2.52 (mg/g) in fresh weight (FW)^6^. Thus, we choose a value of *K*_*B*_ = 2500 (*µ*g/g), which becomes *K*_*B*_ = 100 (*µ*g/cm^2^) after multiplying the conversion factor for petal mass density.

(*σ*) The substrate consists of the proteins, RNA, and nutrients required by viruses to proliferate and spread, and *σ* denotes the rate of logistic growth for this substrate. Since the substrate consists of important nutrients for the tulip as well, it is reasonable to assume that the growth of the substrate is approximate to the growth of the perianth. To estimate this, we used a graph that tracked growth of the mass of the perianth of the flower over time^6^, and another graph that tracked the growth of the petal mass and area over time^2^. Van Damme et al. noted that the flowering period occured between 6-8 weeks, and since they only measured the fresh weight of the perianth every week, there were only three data points to analyze^6^. Azad et al. measured the petal area and petal mass of a tulip, but measured starting from the opening of the flower, rather than from the bud^2^. Orders of magnitude for *σ* ranged from 10^−2^ to 10^−1^ h^−1^. We chose *σ* = 0.75 h^−1^.

(*γ*) The viral decay rate’s cooperative growth was modeled from Moreno et al. ^1^, but it could also be related to the half life of mRNA in the plant *Arabidopsis thaliana* (thale cress). Narsai et al. ^17^ treated suspension cell cultures with specific mRNA inhibitors, and measured a wide range of mRNA half lives, from 12 minutes to over 24 hours. Other sources support 40 minutes to a few hours as an appropriate half life for proteins within organic systems^8^. If we choose a half life of roughly 4.6 hours and rewrite the half life function into an exponential function via *γ* = ln 2/*τ* (where *τ* is the half life), we obtain a decay rate of *γ* = 0.15 h^−1^.

To verify the validity of our choice of parameter values, we can perform a double integral on the petals which returns the mass of the biochemical in units of *µ*g. We can see if the resulting concentration is within expected orders of magnitude by dividing the amount by the weight of a fully grown petal, which is roughly 0.8 to 0.9 g^2^. Realistically, *T* ≈ 10^2^ (*µ*g/g), *B* ≈ 10^3^ (*µ*g/g), and *V* ≈ 10^2^ to 10^3^ (*µ*g/g). If we choose a weight of around 0.8 g, integrating the results obtained from the backward model simulations with an initial width of 4.8 cm (Fig. 3 of the main document) yields values of *T* = 316 (*µ*g/g), *B* = 5181 (*µ*g/g), and *V* = 553 (*µ*g/g). Similarly, integrating the results obtained from the forward model simulations with an initial width of 0.526 cm yields values of *T* = 62 (*µ*g/g), *B* = 1312 (*µ*g/g), and *V* = 128 (*µ*g/g). Most values fall within the desired range; however, the concentration of anthocyanin in the forward model resulted in a lower than expected value, but this seems reasonable as it is expected that the TBV lowers the concentration of pigments in infected petals.

### 2. Supplement 2: Nondimensionalization

Our main model for the TBV, including spatial spread for the anthocyanin *T* (*x, t*), the virus load *V* (*x, t*), and the substrate *B*(*x, t*), is given as

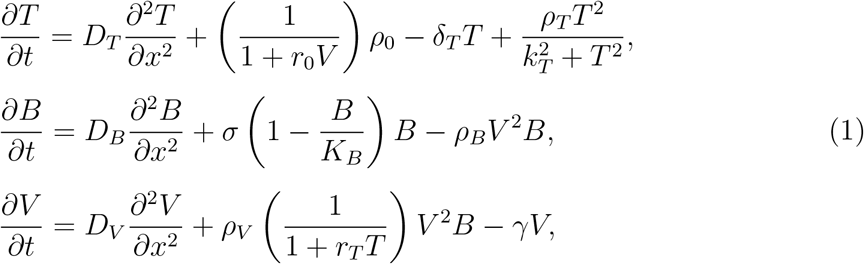

on the domain [0, *L*(*t*)] with homogeneous Neumann boundary conditions

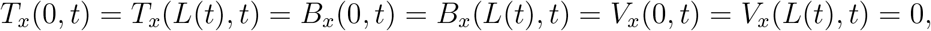

where the index *x* denotes the partial spatial derivative.

To reduce the number of parameters, we nondimensionalize model (1) using the transformations listed in Table 2, and obtain

**Table 2:**
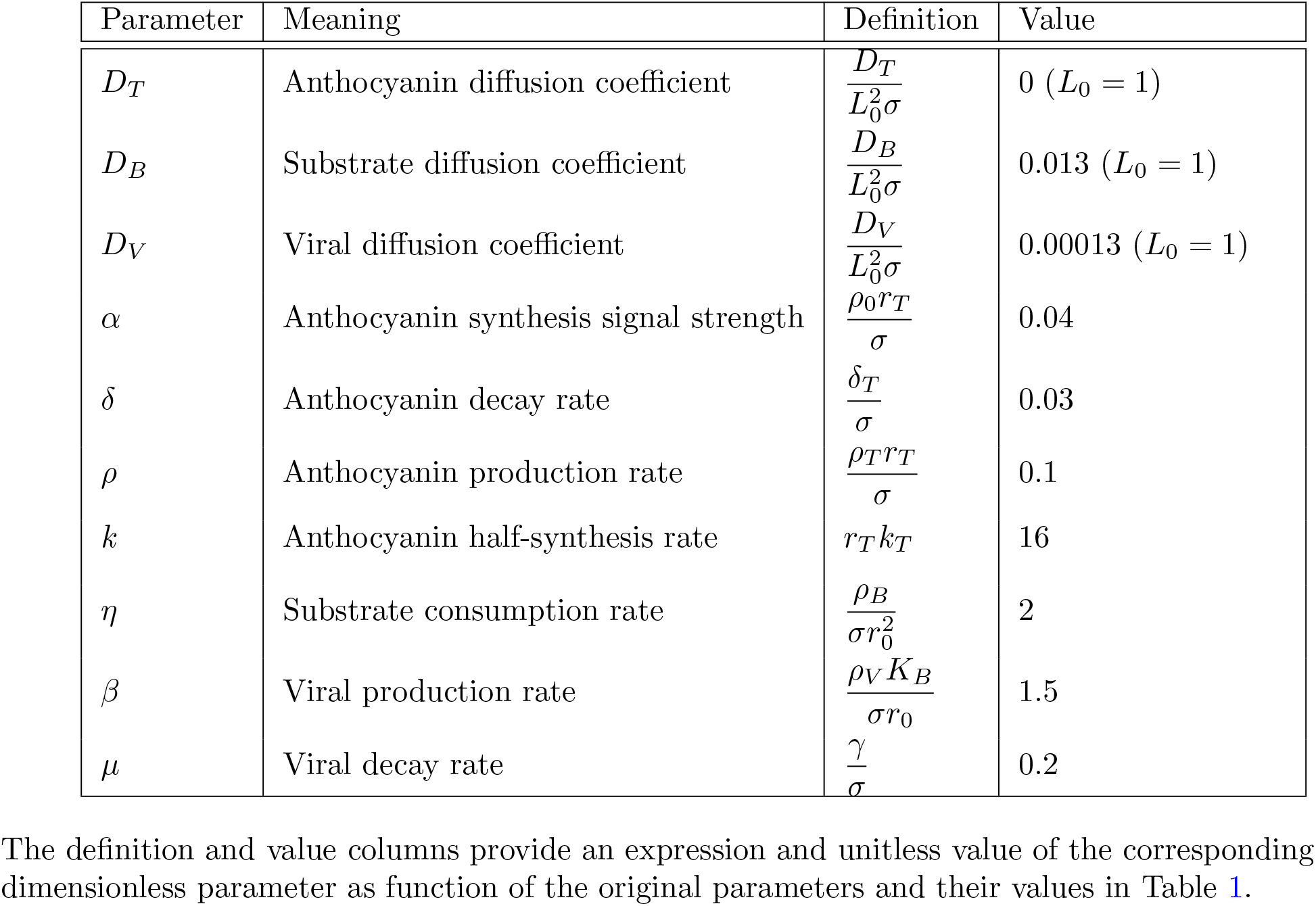
Nondimensionalized parameters of the TBV Model.

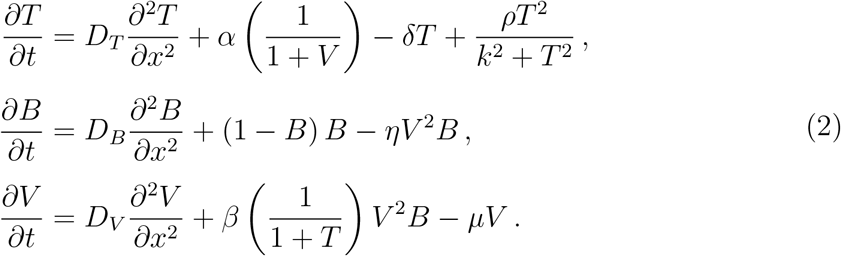

Although the concentrations *T*, *B*, and *V* retain the same symbol as before, they have been multiplied with a scaling factor. Each variable and parameter is now unitless. The nondimensional model (2) is used for a detailed stability and multiscale analysis in Supplement 3.

### 3. Supplement 3: Multiscale Analysis

The parameter values, given in Table 2, for the *T* equation in the nondimensional model (2) are *D*_*T*_ = 0, *α* = 0.04, *δ* = 0.03, *ρ* = 0.1, whereas the parameters for the *B* and *V* equations have values of *D*_*B*_ = 0.013, *η* = 2, *D*_*V*_ = 0.00013, *β* = 1.5, *µ* = 0.2, and a substrate growth rate equal to 1. Hence, the parameters of the *T* equation are about one order of magnitude smaller. We reflect this by introducing a small parameter *ε* to model (2), and obtain

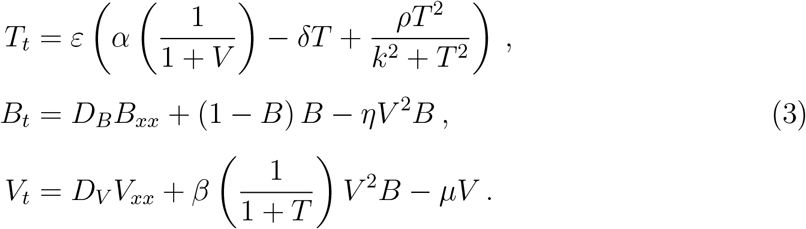

We do not introduce new symbols for the parameters, as for example 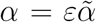, to keep the notation convenient.

#### 3.1 Multiscale Analysis: The Fast System

To leading order in *ε*, we find that the *T* equation simplifies to *T*_*t*_ = 0. Since at early times, when *t* ≈ 0, the amount of anthocyanin can be taken as constant and homogeneous in space, we can assume that *T* will stay constant on the fast time scale. The remaining two equations of system (3) comprise the so called *fast system*, which is given by

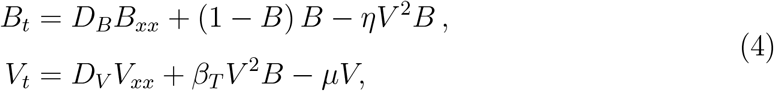

with *β*_*T*_ = *β*(1 + *T*)^−1^. The Jacobian of the kinetic part at a steady state 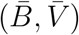 is given as

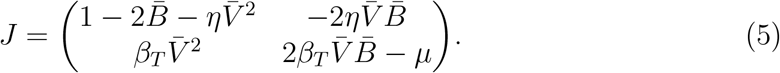

To be able to generate diffusion-driven instabilities, it is necessary for the steady state 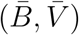 of the fast system (4) to be linearly stable in the absence of diffusion, i.e. we require

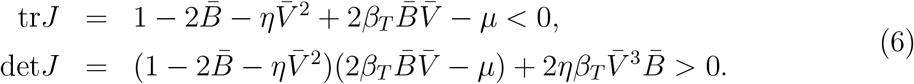

Then, from the well known Turing stability analysis ^7;14^, we can obtain diffusion-driven instability if the Jacobian has one of the two sign patterns

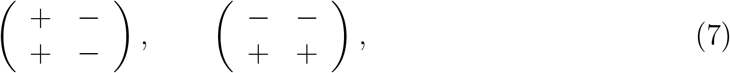

where the left case corresponds to the classical *activator-inhibitor system*, while the right sign pattern corresponds to a *resource-activator system*^16^. Looking at the sign pattern of our Jacobian in (5), we see that for large *η* and small *µ* we might be in the resource-activator case. Let us make this precise.

The homogeneous steady states of system (4) satisfy the two equations

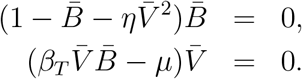

We see that the solution includes 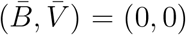 and 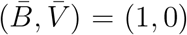. The first point is a saddle point according to the Jacobian (5), while the latter is an asymptotically stable node. Neither of these falls under either sign pattern in (7).

Note that a coexistence steady state 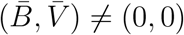 satisfies

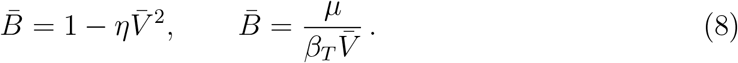

Thus, a nontrivial steady state exists if

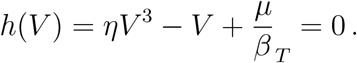

The function *h*(*V*) is a third order polynomial with positive leading order term. We have *h*(0) *>* 0 and *h*^*′*^(*V*) = 0 for two values

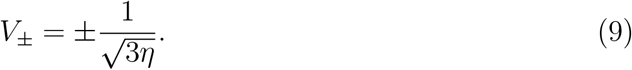

This means we have a local maximum at 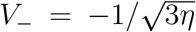 and a local minimum at 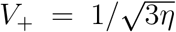. If this latter minimum is below zero, then we have two nontrivial solutions for *h*(*V*) = 0. The value at the minimum is

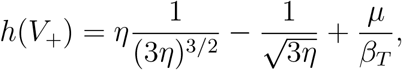

which is less than zero for

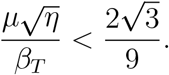

Since *T* ≥ 0 and *β*_*T*_ = *β*(1 + *T*)^−1^, the above condition implies

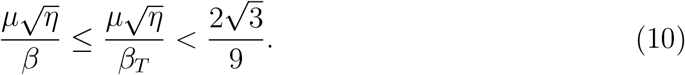

Condition (10) is a necessary conditon for pattern formation, and we assume from here onward that it is satisfied. In this case, we are in the presence of two nonzero steady states,

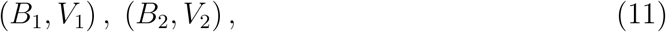

where *h*(*V*_*i*_) = 0 and *B*_*i*_ is given by (8), for *i* = 1, 2.

The Jacobian at these two points becomes

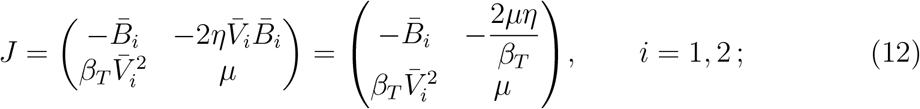

and they both satisfy the sign pattern of the resource-activator case. Next, we need to check which of the two steady states 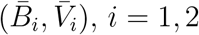, is linearly stable in the absence of diffusion, i.e. satisfies the conditions (6). After substituting 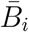 from (8) into the Jacobian, we find that our trace and determinant are

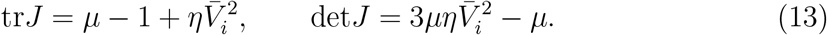

For the steady state 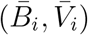 to be linearly stable we require that tr*J <* 0 and det*J >* 0. The determinant condition implies that

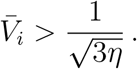

Recall that the minimum of our function *h*(*V*) occurred at this value 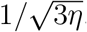. Hence, from the two zeroes of *h*(*V*), the smaller 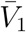 is less than this value and the larger one 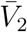 is bigger than this value, i.e.,

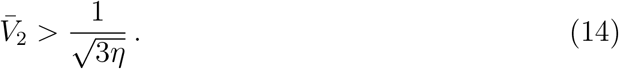

Thus, we have shown that

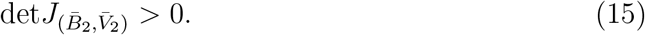

The trace condition tr*J* < 0 for 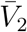 can be rewritten as

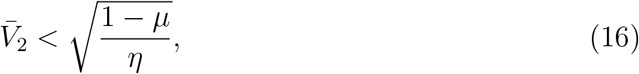

which combined with our previous inequality (14) for 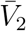 becomes

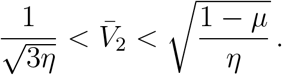

If we use the transitive property, we will have the following necessary condition for the system’s stability without diffusion

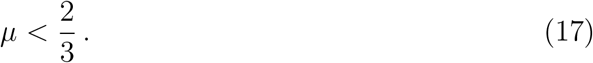

This restriction on the value of *µ* becomes very useful at the time of choosing parameter values.

We summarize these conditions, together with a condition on the diffusion coefficients, in the following Lemma.

##### Lemma 1

*Consider system* (4) *on the interval* [0, *L*] *with homogeneous Neumann boundary conditions*.

1. *If* 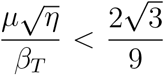, *then there exists a positive homogeneous steady state* 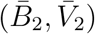 *with the following necessary and sufficient conditions for diffusion-driven instability*
  i. 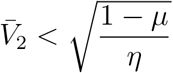,
  ii. 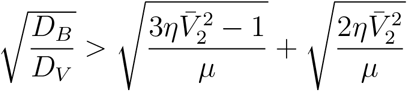,
  iii. *There exist modes* 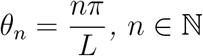 *such that* 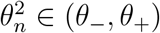, *where*

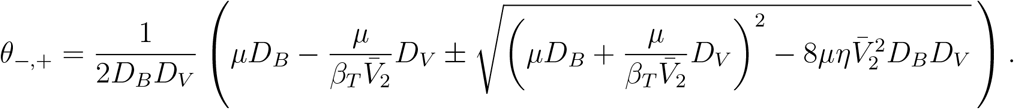 *The modes* 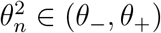 *are linearly unstable for system* (4).
2. *It is convenient to also list necessary conditions for diffusion-driven instability*

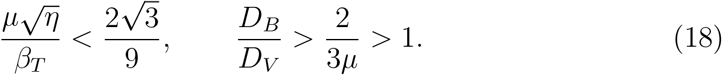

**Proof**.

The existence of the steady state 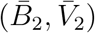 given that 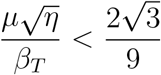 was established in equations (10) and (11).

Denoting the Jacobian at 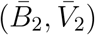 as (see equation (12))

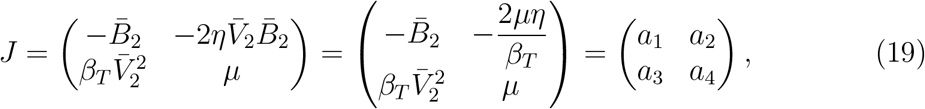

it is known that the necessary and sufficient conditions for diffusion-driven instability in the presence of homogeneous Neumann boundary conditions are given by^7;14^

1. tr *J <* 0,
2. det *J >* 0,
3. 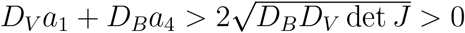.
4. Given the modes 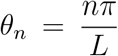, there exists at least one integer *n* > 0 such that 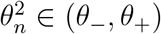 where

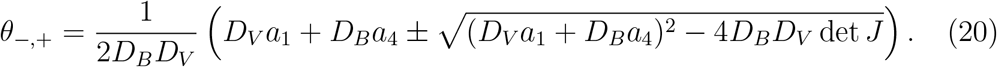

Note that we proved that the second of these conditions is always satisfied (see equation (15)). Also, remember that the trace condition is given by equation (16), or equivalently by condition *i*. in part 1. of the Lemma.

Now, we simplify the third of the above conditions for diffusion-driven instability (inequality condition on the diffusion coefficients) by introducing 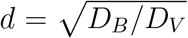, and dividing both sides of the inequality by *d*, to obtain

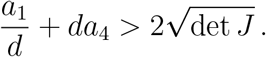

Solving for *d*, using the quadratic equation, yields

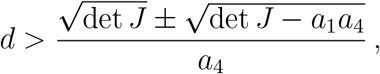

and since 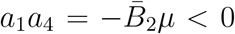, only the positive term is relevant. Using the definition for det *J* = *a*_1_*a*_4_−*a*_2_*a*_3_ and reverting back to the diffusion coefficients provides the following condition for diffusion-driven instability

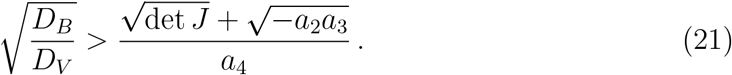

Substitution of expression (13) for the det *J* and the entries of the Jacobian matrix (19) in equation (21) leads to

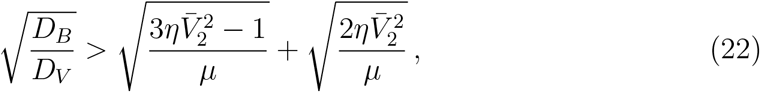

which is condition *ii*. in part 1. of the Lemma.

Note that this last condition guarantees the existence of a nonempty interval (*θ*_−_, *θ*_+_), where *θ*_−_ and *θ*_+_ are given by equation (20). Moreover, considering the definition for det *J* = *a*_1_*a*_4_ − *a*_2_*a*_3_, we can rewrite equation (20) as

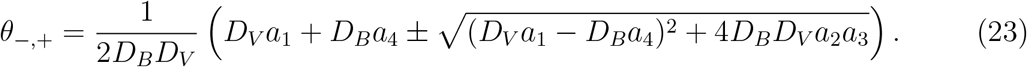

By substituting 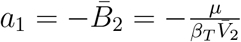(see equation (8)), *a*_4_ = *µ*, and 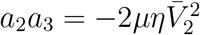 into equation (23), we obtain that

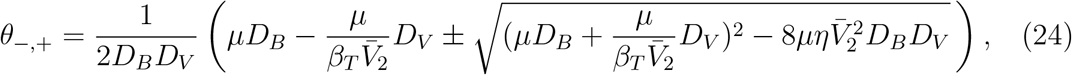

which are precisely the endpoints of the interval (*θ*_−_, *θ*_+_) given by condition *iii*. in part 1 of the Lemma. Thus, the unstable modes are given by

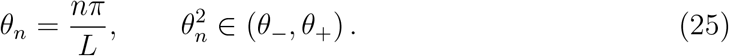

This concludes the proof of the first part of the Lemma.

For the second part of the Lemma, note that the first necessary condition in equation (18) was established already in equation (10). The second necessary condition comes from noticing that both terms on the right hand side of the inequality (22) are positive; thus, for the inequality to hold it is necessary that

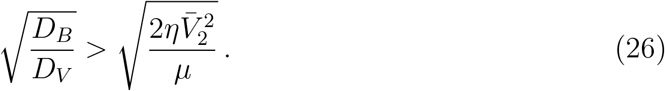

By using the inequality (14) for 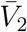 and the necessary condition (17) on *µ* for diffusion-driven instability, equation (26) becomes

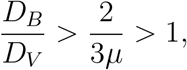

which the second necessary condition in equation (18). □ Note that the condition for the diffusivity ratio to be greater than one is in accordance to the fact that the diffusivity of the substrate is greater than that of the virus. Also notice that, considering equation (10), the necessary conditions (18) for diffusion-driven stability, can be relaxed as the following necessary conditions

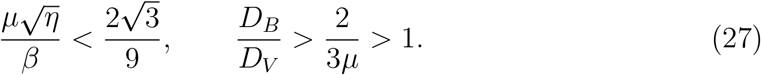

In terms of the dimensional coordinates, these necessary conditions for diffusion-driven instability conditions become

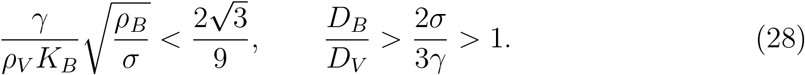

These necessary conditions are helpful to identify parameter values that may lead to diffusion-driven instability.

Consider the unstable modes given by equation (25). As *n* denotes the wavenumber, the shape of the viral load pattern is dominated by the term 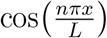, such that for *n* = 2, we would have one local extrema (i.e. one stripe), for *n* = 4 we would expect two stripes, for *n* = 6, three stripes, and so on.

An important feature of reaction-diffusion equations on a growing domain is the factor *L*(*t*)^−2^ that multiplies the diffusion coefficient (see equation (36) in Supplement 4). In addition to changing the criteria for diffusion driven instability, this factor can change the number of stripes at a given time. In general, the larger the domain length *L*(*t*), the larger the modes are, and the greater their range.

#### 3.2. Multiscale Analysis: Slow System

To obtain the corresponding *slow system*, we re-scale time to a long time scale *τ* = *εt* in model (3), and observe that to leading order

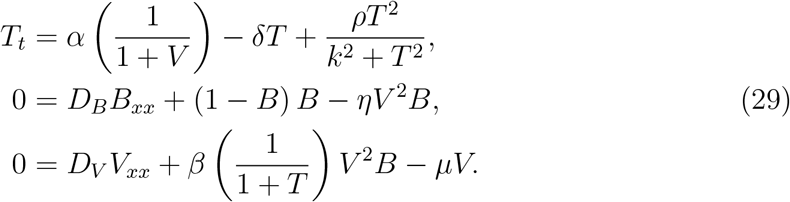

Hence, *B* and *V* satisfy a system of elliptic equations (called the *slow manifold*), and *T* satisfies a Wolpert model of gene activation^13^. The term *α*(1 + *V*)^−1^ is the anthocyanin activation term, which is large if the virus concentration is low and vice versa. In Figure 1, we plot the right hand side of the *T* equation,

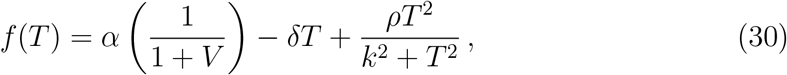

for the two cases of high and low viral load. If the virus concentration is low, *T* converges to a steady state of full anthocyanin expression, while for high *V* we obtain two additional steady states. The lower steady state is also stable and corresponds to anthocyanin inhibition.

**Figure 1:**
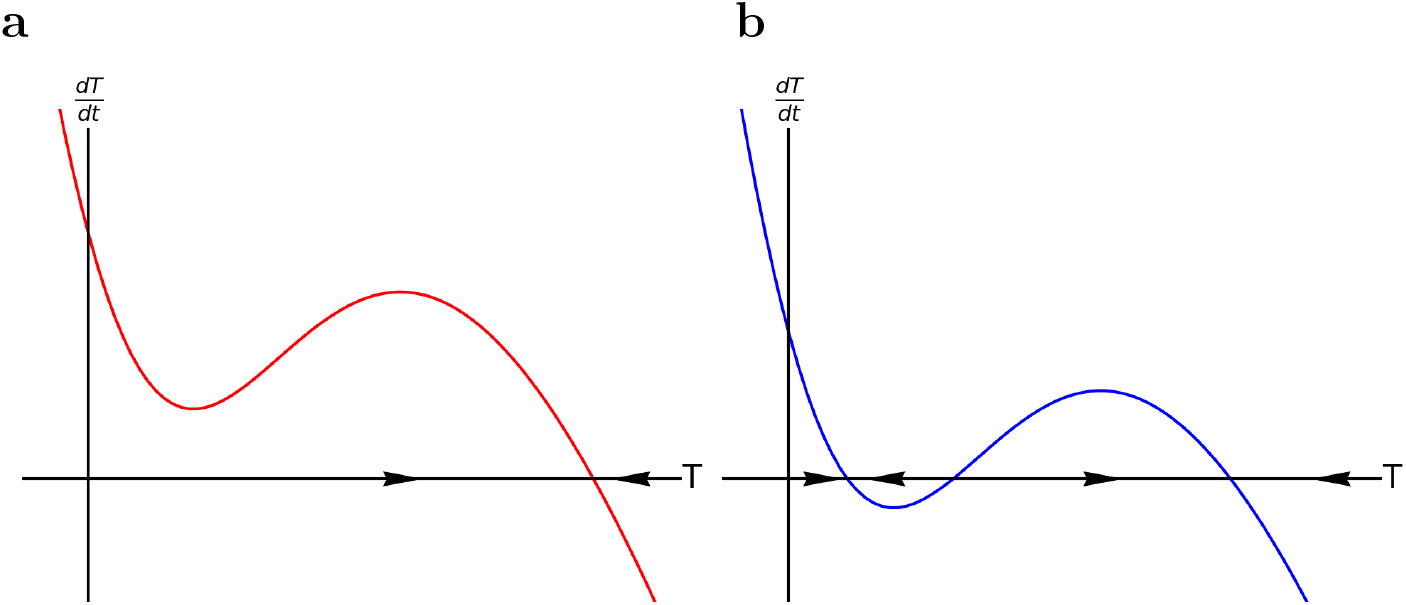
These graphs portray the vector field *f*(*T*) (given by equation (30)) of the Wolpert model with respect to the concentration of anthocyanin *T*. **a** The red curve denotes the rate of change of anthocyanin concentration with a low amount of viruses. There is a single steady state at a high concentration of anthocyanin. **b** The blue curve denotes the rate of change of anthocyanin concentration with a high amount of viruses, weakening the anthocyanin gene signal and creating a new stable state at a much lower concentration of anthocyanin.

### 4. Supplement 4: Reaction-diffusion equations on a growing domain

Reaction-diffusion equations on a growing domain [0, *L*(*t*)] have been thoroughly studied by Crampin et al. ^4^. In this section, we use their methodology for transforming the TBV model (1) on a growing domain [0, *L*(*t*)] into a modified reaction-diffusion system on a fixed domain [0, 1].

The TBV model (1) can be rewritten as the following system

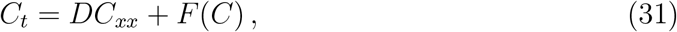

with

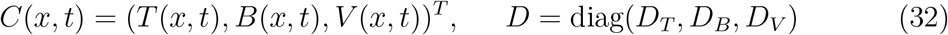

and

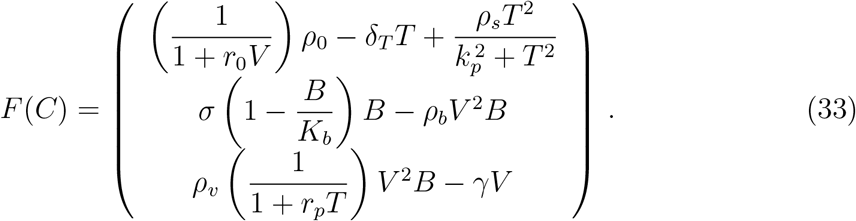

Using the conservation equation and the Reynolds transport theorem, Crampin et al.^4^ showed that the associated evolution equation to a reaction-diffusion system like the one given by equation (31) on a growing domain 0 ≤ *x* ≤ *L*(*t*) is given by

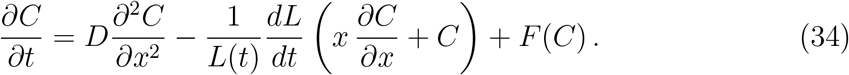

For the purpose of carrying out numerical simulations of the TBV model (31), we transform the spatial variable, as suggested in Crampin et al. ^4^, to a fixed unit interval domain by letting

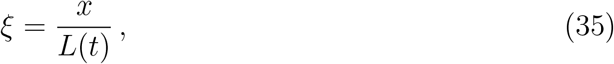

and rewrite (34) as

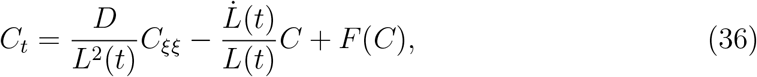

on a fixed unit interval domain 0 ≤ *ξ* ≤ 1.

This allows us to solve numerically the TBV model (31) on the fixed domain 0 ≤ *ξ* ≤ 1, and then use the spatial transformation given by equation (35) to find the numerical solution on the growing domain 0 ≤ *x* ≤ *L*(*t*).

### 5. Supplement 5: Estimating the arc length

This section describes the methodology used to estimate the lengths of the arcs perpendicular to the veins observed on the light image of the tulip petal of the variety *“Tulipa Seadov”*. First, we assume that the arcs perpendicular to the veins observed on the light image of the tulip petal are cells of approximate equal age, and they form arcs of a circle. This reduces the task of measuring the arc lengths to calculating lengths of circular arcs.

Let us see how the lengths of these circular arcs can be calculated only in terms of the half length *a* of the chord with ends given by the intersection of the arcs with the edge of the petal and the distance *b*, along the proximodistal axis of the petal, from the chord to the circular arc (see Figure 2).

First, note that this geometrical setup defines two similar triangles enclosed in a circle with the arc whose length we want to calculate (green circular arc in Figure 2). Thus, equating the ratios of the corresponding sides of these similar triangles provides the radius of the circle,

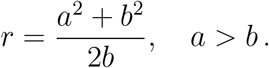

Having the radius *r* of the circle in terms of *a* and *b*, and knowing that the circular arc length is given by

**Figure 2:**
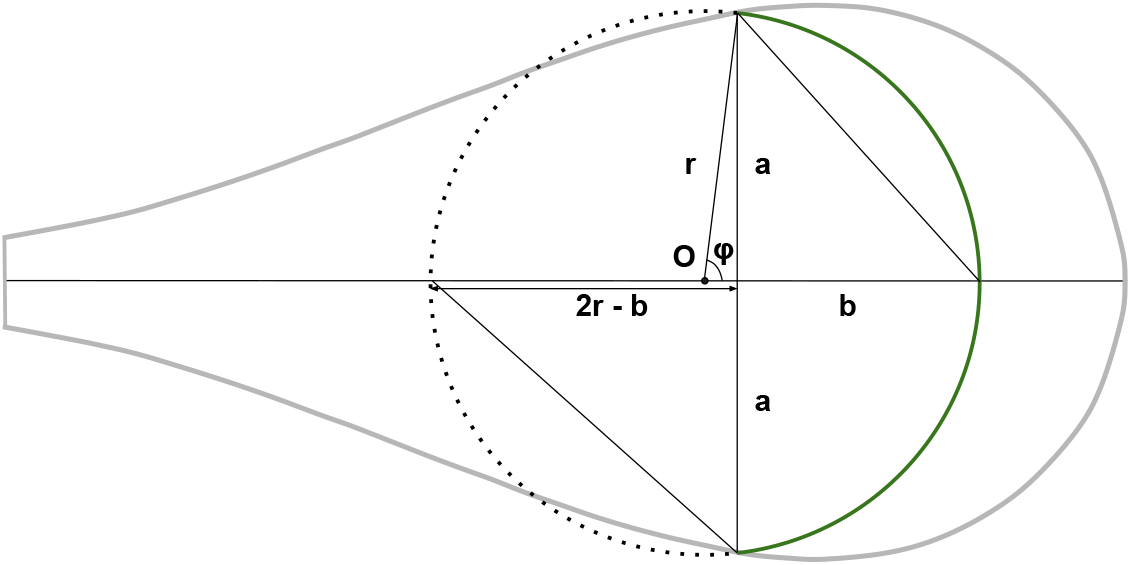
A band of equally aged cells (in green) can be interpreted as part of a circle. The radius and arc length of this band can be determined purely from the half chord length *a* and chord height *b*.

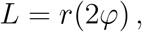

where 2*φ* denotes the central angle of the arc (see Figure 2), leaves us only with the task of expressing *φ* in terms of *a* and *b*. For this, notice that

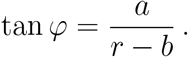

Thus,

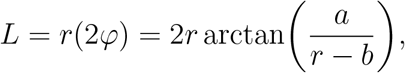

or in terms of only *a* and *b*,

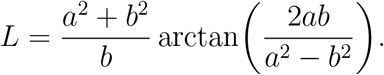

Note that when *b* approaches 0, *L* approaches 2*a* as expected, and when *b* approaches *a, L* approaches *πa*, also as expected.

Having the values for *a* and *b* allows us not only to assign a circular geometry to the arcs that are perpendicular to the veins of the petal and represent the equally aged cells but also to estimate the lengths of the arcs, data that is used to describe the growth and curvature of a tulip petal.

### 6. Supplement 6: Computer Code

All simulations were run on the programming language *Python*. The packages *numpy, matplotlib*, and *scipy* were used. A study on Turing patterns by Iain Barr^3^ served as the foundation of our code, but while he uses Euler forward methods, our PDEs were solved using the Crank-Nicholson method to form a tri-diagonal matrix to make the numerics more stable, followed by a Thomas algorithm. The code is available on Github ^1^.

## Footnotes

1 https://github.com/aidanawong/reaction-diffusion-growing-domain

## Notes

### Competing Interest Statement

The authors have declared no competing interest.

## References

[1] Agu-Barragan A, Ochoa-Alejo N (2014) Virus-induced silencing of myb and wd40 transcription factor genes affects the accumulation of anthocyanins in chilli pepper fruit. Biologia Plantarum 58:567–574

[2] Andreu-Moreno I, Bou JV, Sanjuán R (2020) Cooperative nature of viral replication. Science Advances 6(49). 10.1126/sciadv.abd4942

[3] Artist UD (17th Century) Great tulip book: Kamelot van wena. URL https://www.nortonsimon.org/art/detail/M.1974.08.086.D, [Online; accessed November 23, 2023]

[4] Artist UD (17th Century) Great tulip book: Witte mervelije. URL https://www.nortonsimon.org/art/detail/M.1974.08.149.D, [Online; accessed November 23, 2023]

[5] Ben-Ari ET (1999) The silence of the genes: In a game of evolutionary one-upmanship, plants and viruses wrangle over gene silencing. BioScience 49(6):432–437. 10.2307/1313550, URL https://doi.org/doi.org/10.2307/1313550, https://arxiv.org/abs/https://academic.oup.com/bioscience/article-pdf/49/6/432/19403079/49-6-432.pdfhttps://academic.oup.com/bioscience/article-pdf/49/6/432/19403079/49-6-432.pdf

[6] Crampin EJ, Gaffney EA, Maini PK (1999) Reaction and diffusion on growing domains scenarios for robust pattern formation. Bulletin of Mathematical Biology 61(6):1093–1120. 10.1006/bulm.1999.0131, URL https://www.sciencedirect.com/science/article/pii/S0092824099901313

[7] Ding B, Patterson EL, Holalu SV, et al (2020) Two myb proteins in a self-organizing activator-inhibitor system produce spotted pigmentation patterns. Current Biology 30:802–814.e8

[8] Dubos R (1959) Tulipomania and the Benevolent Virus. Perspectives in Virology. John Wiley and Sons, New York

[9] Espley RV, Brendolise C, Chagné D, et al (2009) Multiple repeats of a promoter segment causes transcription factor autoregulation in red apples. The Plant Cell 21(1):168–183. 10.1105/tpc.108.059329, URL https://doi.org/10.1105/tpc.108.059329, https://arxiv.org/abs/https://academic.oup.com/plcell/article-pdf/21/1/168/36917077/plcellv211168.pdfhttps://academic.oup.com/plcell/article-pdf/21/1/168/36917077/plcellv211168.pdf

[10] Gierer A, Meinhardt H (1972) A theory of biological pattern formation. Kybernetik 12:30–39

[11] Green JBA, Sharpe J (2015) Positional information and reaction-diffusion: two big ideas in developmental biology combine. Development 142(7):1203–1211

[12] Inukai T, Kim H, Matsunaga W, et al (2022) Battle for control of anthocyanin biosynthesis in two brassicaceae species infected with turnip mosaic virus. Journal of Experimental Botany 74(5):1659–1674. 10.1093/jxb/erac502, URL https://doi.org/10.1093/jxb/erac502, https://arxiv.org/abs/https://academic.oup.com/jxb/article-pdf/74/5/1659/49646719/erac502.pdfhttps://academic.oup.com/jxb/article-pdf/74/5/1659/49646719/erac502.pdf

[13] Jorgensen RA (1995) Cosuppression, flower color patterns, and metastable gene expression states. Science 268(5211):686–691. 10.1126/science.268.5211.686

[14] Leipziger D (2013) Flower Map. Finishing Line Press, Georgetown, KY

[15] Lerner RB (2005) Forcing bulbs for indoor bloom

[16] Leshchiner A, Solovyev A, Morozov S, et al (2006) A minimal region in the ntpase/helicase domain of the tgbp1 plant virus movement protein is responsible for atpase activity and cooperative rna binding. The Journal of general virology 87:3087–95. 10.1099/vir.0.81971-0

[17] Lesnaw J, Ghabrial S (2000) Tulip breaking past, present, and future. Plant Disease - PLANT DIS 84:1052–1060. 10.1094/PDIS.2000.84.10.1052

[18] Lewis J, Slack J, Wolpert L (1977) Thresholds in development. Journal of Theoretical Biology 65(3):579–590. 10.1016/0022-5193(77)90216-8, URL https://www.sciencedirect.com/science/article/pii/0022519377902168

[19] Luo QJ, Mittal A, Jia F, et al (2012) An autoregulatory feedback loop involving pap1 and tas4 in response to sugars in arabidopsis. Plant molecular biology 80:117–129. 10.1007/s11103-011-9778-9

[20] Maini PK, Woolley TE (2019) The turing model for biological pattern formation. In: Bianchi A, Hillen T, Lewis MA, et al (eds) The Dynamics of Biological Systems. Springer International Publishing, chap 7, p 189–204, 10.1007/978-3-030-22583-4_7

[21] Miura T (2013) Turing and wolpert work together during limb development. Science Signaling 6(270):pe14

[22] Moe R, Wickstrøm A (1973) The effect of storage temperature on shoot growth, flowering, and carbohydrate metabolism in tulip bulbs. Physiologia Plantarum 28(1):81–87. 10.1111/j.1399-3054.1973.tb01155.x

[23] Moelling K (2016) Tulipomania - the first financial crisis by viruses. Revue Roumaine de Chimie 61:637–645

[24] Moelling K (2017) Viruses: More Friends than Foes. World Scientific, Berlin

[25] Mort R, Ross R, Hainey K, et al (2016) Reconciling diverse mammalian pigmentation patterns with a fundamental mathematical model. Nature Communications 7:10288

[26] Painter K, Othmer H, Maini P (1999) Stripe formation in juvenile pomacanthus via chemotactic response to a reaction-diffusion mechanism. Proceedings of the National Academy of Sciences 96:5549–5554

[27] Petroni K, Tonelli C (2011) Recent advances on the regulation of anthocyanin synthesis in repro-ductive organs. Plant science 181:219–229. 10.1016/j.plantsci.2011.05.009

[28] Qian H, Murray J (2001) A simple method of parameter space determination for diffusion-driven instability with three species. Applied Mathematics Letters 14(4):405–411. 10.1016/S0893-9659(00)00169-5, URL https://www.sciencedirect.com/science/article/pii/S0893965900001695

[29] Saha S, Lõhmus A, Dutta P, et al (2022) Interplay of hcpro and cp in the regulation of potato virus a rna expression and encapsidation. Viruses (Basel) 14(6). 10.3390/v14061233

[30] Segredo-Otero E, Sanjuán R (2022) Cooperative virus-virus interactions: An evolutionary perspective. BioDesign Research

[31] Sosnov V, Ulrychová M (1972) Tobacco mosaic virus reproduction in plants with an increased anthocyanin content induced by phosphorus deficiency. Biologia Plantarum 14:133–139

[32] The Mount Vernon Ladies’ Association (2020) Absalon tulip. URL https://www.mountvernon.org/the-estate-gardens/gardens-landscapes/plant-finder/item/absalon-tulip/, [Online; accessed October 31, 2023]

[33] Thompson EA (2007) The tulipmania: Fact or artifact? Public Choice 130:99–114

[34] Tian J, Zhang J, Han Zy, et al (2017) Mcmyb12 transcription factors co-regulate proantho-cyanidin and anthocyanin biosynthesis in malus crabapple. Scientific Reports 7:43715. 10.1038/srep43715

[35] Turing A (1952) The chemical basis of morphogenesis. Bulletin of Mathematical Biology 52:153–197

[36] Wang Y, Chen L, Yang Q, et al (2022) New insight into the pigment composition and molecular mechanism of flower coloration in tulip (tulipa gesneriana l.) cultivars with various petal colors. Plant Science 317:111193. 10.1016/j.plantsci.2022.111193, URL https://www.sciencedirect.com/science/article/pii/S0168945222000176

[37] wu Yuan Y, Sagawa JM, Frost LA, et al (2014) Transcriptional control of floral anthocyanin pigmentation in monkeyflowers (mimulus). The New phytologist 204 4:1013–27

[38] Zaitlin M (1998) The Discovery of the Causal Agent of the Tobacco Mosaic Disease, chap 7, pp 105–110. 10.1142/9789812817563\_0007, URL https://www.worldscientific.com/doi/abs/10.1142/9789812817563\_0007, https://www.worldscientific.com/doi/pdf/10.1142/9789812817563\_0007

## References

[1] Andreu-Moreno I, Bou JV, Sanjuán R (2020) Cooperative nature of viral replication. Science Advances 6(49). 10.1126/sciadv.abd4942

[2] Azad AK, Hanawa R, Ishikawa T, et al (2013) Expression profiles of aquaporin homologues and petal movement during petal development in tulipa gesneriana. Physiologia plantarum 148 3:397–407

[3] Barr I (2017) Turing patterns. URL https://www.degeneratestate.org/posts/2017/May/05/turing-patterns/

[4] Crampin EJ, Gaffney EA, Maini PK (1999) Reaction and diffusion on growing domains scenarios for robust pattern formation. Bulletin of Mathematical Biology 61(6):1093–1120. 10.1006/bulm.1999.0131, URL https://www.sciencedirect.com/science/article/pii/S0092824099901313

[5] Cuntz M, Ogée J, Farquhar GD, et al (2007) Modelling advection and diffusion of water isotopologues in leaves. Plant, Cell & Environment 30(8):892–909. 10.1111/j.1365-3040.2007.01676.x

[6] Damme EJMV, Peumans WJ (1989) Developmental changes and tissue distribution of lectin in tulipa. Planta 178:10–18

[7] Edelstein-Keshet L (2005) Mathematical Models in Biology. Society for Industrial and Applied Mathematics, 10.1137/1.9780898719147, URL https://epubs.siam.org/doi/abs/10.1137/1.9780898719147, https://epubs.siam.org/doi/pdf/10.1137/1.9780898719147

[8] Fernandez-Rodriguez J, Voigt CA (2016) Post-translational control of genetic circuits using Potyvirus proteases. Nucleic Acids Research 44(13):6493–6502. https://doi.org/10.1093/nar/gkw537, URL 10.1093/nar/gkw537, https://academic.oup.com/nar/article-pdf/44/13/6493/17437431/gkw537.pdf

[9] Guo X, Fu X, Li X, et al (2022) Effect of flavonoid dynamic changes on flower coloration of tulipa gesneiana ‘queen of night’ during flower development. Horticulturae 8(6). 10.3390/horticulturae8060510, URL https://www.mdpi.com/2311-7524/8/6/510

[10] Hagiladi A, Bourque D, Wildman S (1975) Rates of rod accumulation and viral rna synthesis during early and late stages of tobacco mosaic virus infection in young, expanding tobacco leaves. Virology 63(1):123–129. 10.1016/0042-6822(75)90377-3, URL https://www.sciencedirect.com/science/article/pii/0042682275903773

[11] Hayashi K, Mori M, Knox YM, et al (2003) Anti influenza virus activity of a red-fleshed potato anthocyanin. Food Science and Technology Research 9(3):242–244. 10.3136/fstr.9.242

[12] Lellig C, Wagner J, Hempelmann R, et al (2004) Self-diffusion of rodlike and spherical particles in a matrix of charged colloidal spheres: A comparison between fluorescence recovery after photobleaching and fluorescence correlation spectroscopy. The Journal of Chemical Physics 121:7022–9. 10.1063/1.1791631

[13] Lewis J, Slack J, Wolpert L (1977) Thresholds in development. Journal of Theoretical Biology 65(3):579–590. 10.1016/0022-5193(77)90216-8, URL https://www.sciencedirect.com/science/article/pii/0022519377902168

[14] Maini PK, Woolley TE (2019) The Turing model for biological pattern formation. In: Bianchi A, Hillen T, Lewis MA, et al (eds) The Dynamics of Biological Systems. Springer International Publishing, chap 7, p 189–204, 10.1007/978-3-030-22583-4_7

[15] Mavituna F, Park J, Gardner D (1987) Determination of the effective diffusion coefficient of glucose in callus tissue. The Chemical Engineering Journal 34(1):B1–B5. 10.1016/0300-9467(87)85008-6

[16] Meinhardt H (2008) Models of biological pattern formation: From elementary steps to the organization of embryonic axes. Current topics in developmental biology 81:1–63. 10.1016/S0070-2153(07)81001-5

[17] Narsai R, Howell K, Millar A, et al (2007) Genome-wide analysis of mrna decay rates and their determinants. The Plant Cell 19:3418–36. 10.1105/tpc.107.055046

[18] Paine PL, Moore LC, Horowitz SB (1975) Nuclear envelope permeability. Nature 254:109–114. 10.1038/254109a0

[19] Rutschow HL, Baskin TI, Kramer EM (2011) Regulation of solute flux through plasmodesmata in the root meristem. Plant Physiology 155:1817–1826

[20] Sano Y (1987) Translational diffusion coefficient of tobacco mosaic virus particles. Journal of General Virology 68(9):2439–2442. 10.1099/0022-1317-68-9-2439, URL https://www.microbiologyresearch.org/content/journal/jgv/10.1099/0022-1317-68-9-2439

[21] Santos N, Castanho M (1996) Teaching light scattering spectroscopy: the dimension and shape of tobacco mosaic virus. Biophysical Journal 71(3):1641–1650. 10.1016/S0006-3495(96)79369-4, URL https://www.sciencedirect.com/science/article/pii/S0006349596793694

[22] Tan S, Ebrahimi A, Langrish T (2018) Preparation of core-shell microspheres of lactose with flower-like morphology and tailored porosity. Powder Technology 325:309–315. 10.1016/j.powtec.2017.11.028, URL https://www.sciencedirect.com/science/article/pii/S0032591017308951

[23] Tian J, Zhang J, Han Zy, et al (2017) Mcmyb12 transcription factors co-regulate proanthocyanidin and anthocyanin biosynthesis in malus crabapple. Scientific Reports 7:43715. 10.1038/srep43715

[24] Weiss D (2000) Regulation of flower pigmentation and growth: multiple signaling pathways control anthocyanin synthesis in expanding petals. Physiologia plantarum 110(2):152–157

